# LATTE for locus-specific quantification of transposable element expression across species

**DOI:** 10.64898/2026.03.28.714964

**Authors:** Jiayi He, Chen Peng, Yuelang Zhang, Zhengguang Wang, Haihan Zhang, Lingzhao Fang, Pengju Zhao

## Abstract

Transposable elements (TEs) are pivotal drivers of eukaryotic genome evolution and phenotypic diversity. However, their functional contributions to complex traits remain largely obscured by expression quantification challenges arising from high sequence homology and multi-mapping ambiguities. Here, we present **LATTE**, an efficient computational framework for defining and quantifying TE expression at locus-specific resolution by leveraging an innovative multi-indicator Expectation-Maximization (EM) algorithm. Extensive benchmarking against simulated datasets demonstrated that LATTE significantly outperformed existing state-of-the-art tools, achieving an accuracy of 0.998 at the subfamily level and 0.839 at the locus-specific level. Applying LATTE to 813 RNA-seq datasets across humans, cattle, and chickens, we quantified expression profiles of 2,703 TEs, followed by TE-expression quantitative trait loci (TE-eQTL) mapping. The colocalization rates between TE-eQTL and host gene-eQTL was low, revealing a distinct regulatory landscape of TE expression. This decoupled correlation between TEs and host genes are likely mediated by the differential expression of alternative transcripts. Through integrated TE-eQTL and genome-wide association studies on 3,746 complex traits across three species, we demonstrated that TEs constitute 204 (8.7%) additional associations with complex traits beyond gene-eQTL. More specifically, the Sjögren’s syndrome-associated variant rs10954213 acts as a TE-eQTL that shifts the splicing landscape of *IRF5*, upregulating TE-containing transcripts while simultaneously suppressing canonical ones. Collectively, LATTE provides an efficient framework for studying TE expression across species, and our findings highlight the key role of TEs in understanding the genetic architecture of complex phenotypes.

## Introduction

Transposable elements (TEs) are DNA sequences capable of mobilizing within a genome^1^. Based on their transposition mechanisms, terminal repeats, and sequence lengths, TEs are generally classified into short interspersed nuclear elements (SINEs), long interspersed nuclear elements (LINEs), long terminal repeat (LTR) retrotransposons, and DNA transposons^2,3^. These classes are further subdivided into thousands of TE families and subfamilies based on sequence homology^2^. As major components of eukaryotic genomes, TEs play crucial roles in genome evolution and diversity. For instance, TEs constitute approximately 49.1% of the human genome^4^, 46.2% of the cattle genome, and 10.4% of the chicken genome^5^. Notably, the abundance and composition of TEs vary significantly across species: the human genome is dominated by SINE/Alu^4^, the cattle genome by LINE/L1, and the chicken genome by LINE/CR1^5^.

TEs have been extensively documented to be associated with diverse genomic functions^6,7^, and ultimately, complex phenotypes and diseases^8,9^. For instance, high expression of endogenous retroviruses (ERVs), a TE family within the LTR class^10^, is associated with immune deficiencies in patients with systemic lupus erythematosus (SLE) or cancer^11–14^. Furthermore, while some studies have investigated TE expression and its underlying genetic control^15^, such evidence has been limited to the TE subfamily level due to the lack of locus-specific genomic coordinates. Consequently, major international initiatives, such as the human Genotype-Tissue Expression (GTEx) project^16^ and the FarmGTEx project^17–20^, have yet to fully explore the genetic regulation of TEs and their subsequent impacts on complex traits. Therefore, it is of great interest to systematically investigate the spatiotemporal expression patterns of TEs and how they are genetically regulated to influence higher-order phenotypes across a wide range of species.

Due to their mobility and homology, TEs exhibit wide interspersion and high sequence similarity, both within TE families and between TEs and exogenous viruses, throughout the genome (**Figure 1A**). This creates significant challenges in quantifying TE expression from short-read RNA-seq, which is the dominated transcriptome data in public domain (**Figure 1B**): (1) TE-derived reads are often aligned to multiple genomic locations^21^, which complicates locus-specific quantification; (2) many reads map to regions spanning exon-TE junctions^22^ or intergenic regions, meaning that incomplete or inaccurate genomic coordinates can lead to the incorrect identification of TE exonization boundaries, thereby affecting functional annotation; and (3) sequencing error, imperfect alignment, or exogenous interference can introduce mismatches between reads and the reference genome^23^, leading to substantial bias in TE expression estimates. Although there are multiple existing tools for quantification of TE expression ^24–28^, there remains limitations particularly in areas such as cross-species availability, the ability to detect comprehensive profiling of TEs, locus-specific quantification, and the identification of interference (**Figure 1C**).

**Figure 1.**
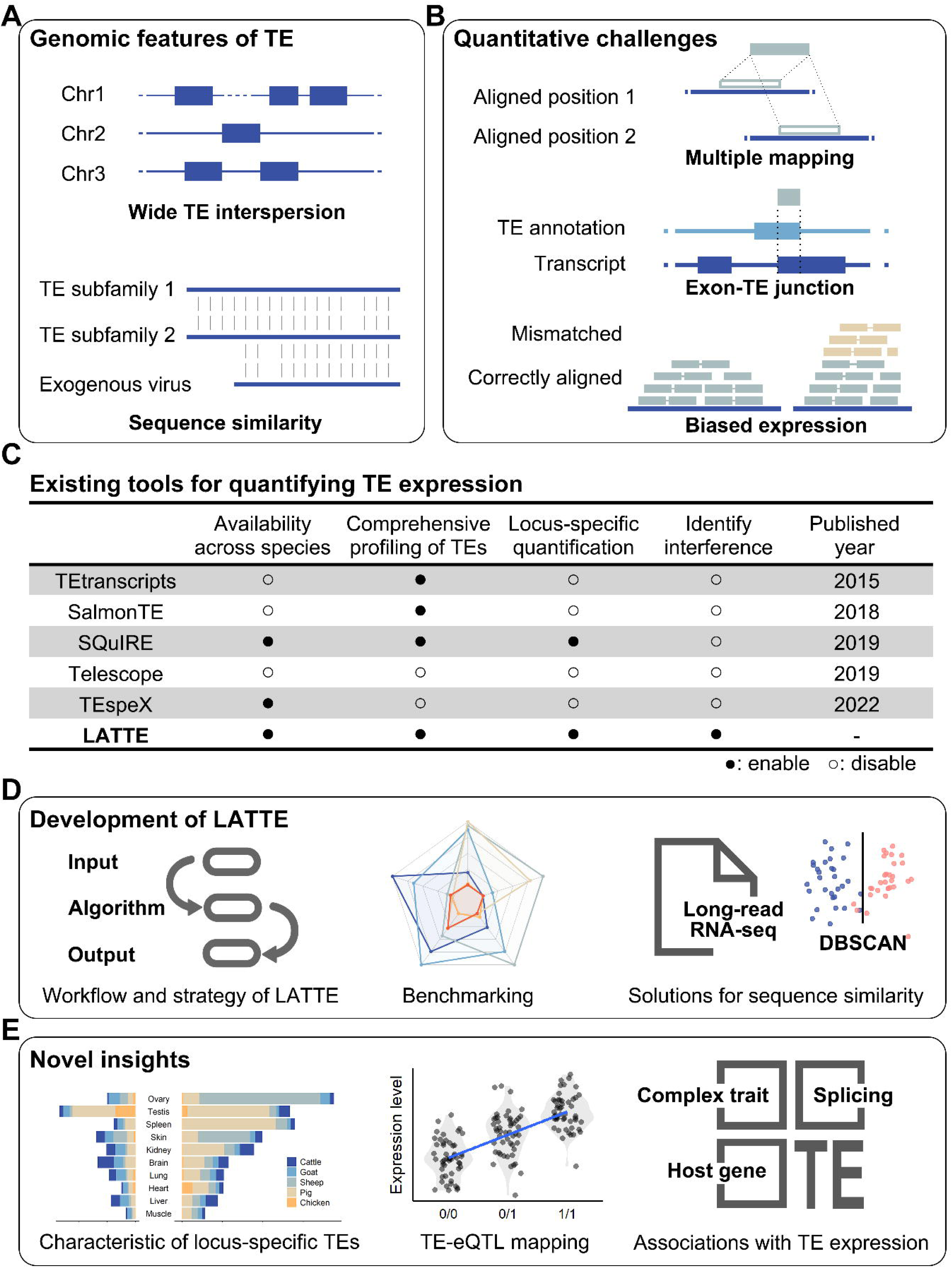
Background, development, and insights of LATTE. (A) Two genomic features of TEs arising from their mobility and homology: wide interspersion and sequence similarity. (B) Three major quantitative challenges for short-read RNA-seq, including multi-mapping, ambiguous genomic coordinates of TE expression, and interference from mismatching or exogenous viruses. (C) Benchmark of basic capabilities of existing TE expression quantification tools. (D) Development of LATTE: the study details its improvements, usage, strategies, benchmarking, and innovative solutions for TE homology. (E) Biological insights into TE expression facilitated by LATTE, including tissues-specific expression patterns, genetic regulation, and associations with biological processes and complex traits.

To bridge this gap, we developed **LATTE** (https://github.com/PengjuZ/LATTE), an efficient computational framework designed for quantifying TE expression at locus-specific resolution. Leveraging an innovative multi-indicator EM algorithm that incorporates subfamily identity, base coverage, and genomic annotation, LATTE accurately resolved mapping ambiguities (**Figure 1D**). As a core tool of the FarmGTEx project^17^, the framework integrated long-read RNA-seq data (*n*=29) from five farmed species to curate high-confidence active TE libraries and employed machine learning-based denoising to mitigate homology-driven biases. We then applied LATTE to study TE expression pattern across tissues by analyzing 250 RNA-seq in five species. Furthermore, we conducted a comprehensive cross-species investigation into the genetic basis of TE expression by analyzing 813 short-read RNA-seq datasets of blood with paired whole-genome sequence (WGS) data in humans (*n*=451), cattle (*n*=170), and chickens (*n*=192) (**Figure 1E**). Our findings revealed that TEs were regulated by independent *cis*-regulatory variants with profound tissue-specificity, representing a functional layer largely decoupled from host genes. Most notably, through transcriptome-wide association studies (TWAS) of 3,746 complex traits across three species, we demonstrated that TEs constitute 204 (8.7%) exclusive associations with complex traits beyond gene-eQTL. Collectively, this study provides an efficient computational framework for high-resolution TE quantification and redefines TEs as active, independent components of the genetic architecture underlying complex phenotypic variation.

## Results

### LATTE enables high-resolution quantification of TE expression via a multi-indicator EM strategy

LATTE introduces four key improvements to address the current limitations in TE expression research mentioned above (**Figure 2A**): (1) an innovative reassignment strategy that enabled more precise quantification of ambiguous reads; (2) read-specific resolution that provided exact coordinates for exon-TE junctions—a level of detail currently unattainable by existing tools; (3) broad species applicability, extending robust performance beyond model organisms to a diverse range of non-model species; and (4) integrated denoising solutions that enhanced the assessment accuracy of homologous TEs.

**Figure 2.**
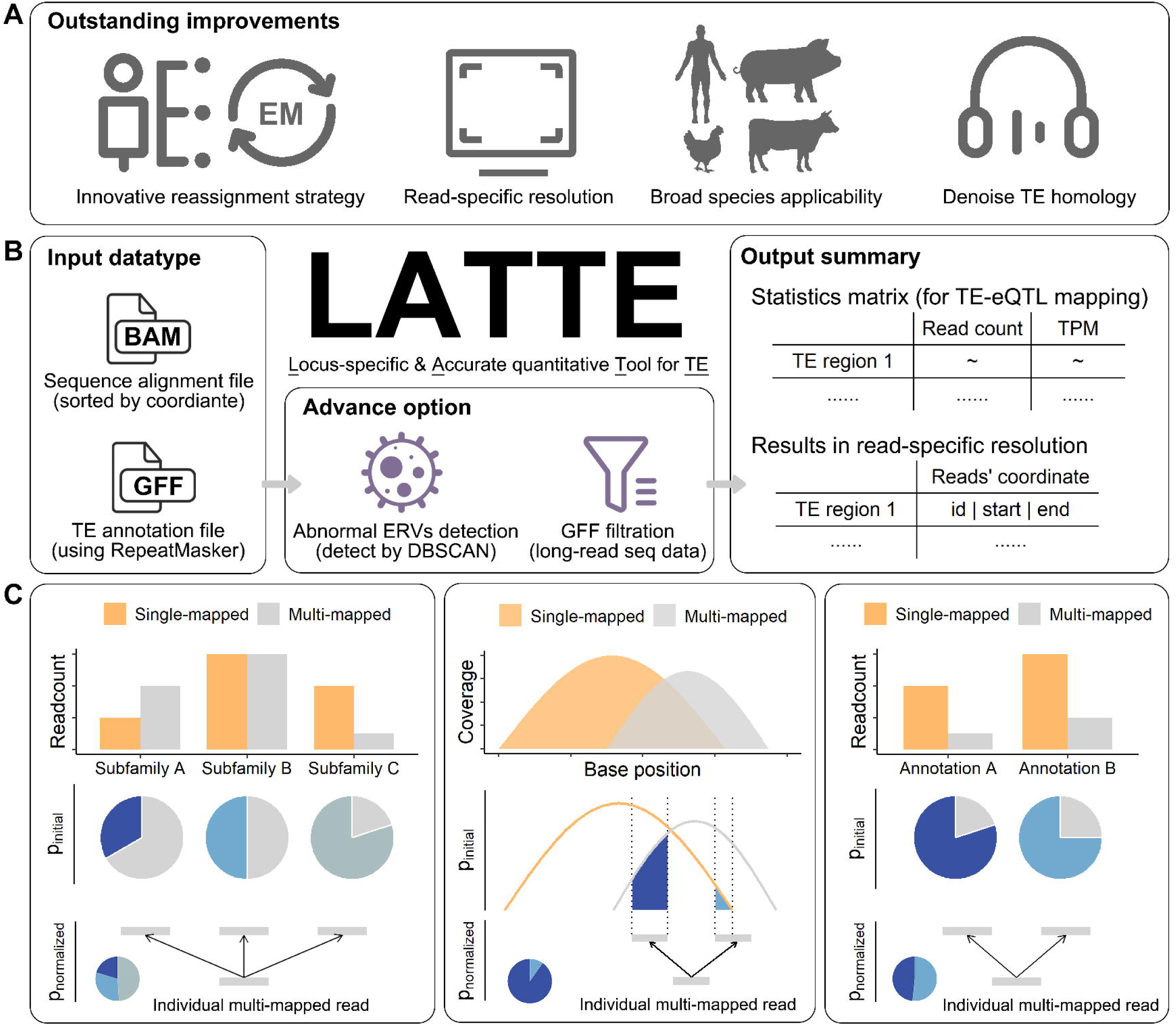
Schematic overview of the LATTE workflow. (A) Four key improvements of LATTE, including the innovative EM-based reassignment strategy, read-specific resolution, broad species applicability, and denoising solutions for TE homology. (B) The end-to-end LATTE analytical pipeline, illustrating input data types, advanced options, and output files. (C) Implementation of the EM algorithm in LATTE, depicting three iterative modules, that calculate probabilities at the TE subfamily, base coverage, and annotated locus levels to assign multi-mapped reads.

LATTE was implemented in Shell and Python, and tested with the following versions of software and packages: Glibc (v2.17), SAMtools (v1.17)^29^, BEDTools (v2.31.1)^30^, Python (v3.11), NumPy (v2.1.0)^31^, Pandas (v2.2.2), and Scikit-learn (v1.5.1)^32^. This tool is designed for UNIX environments and requires no formal installation. As a lightweight framework with minimal constraints, LATTE executes its analysis via a single-line command (see GitHub README for details). Users only need to provide two input files: a coordinate-sorted BAM file of RNA-seq data and a corresponding TE annotation file in GFF format (**Figure 2B**). The quantification process is then executed automatically, typically yielding two primary outputs: a comprehensive record of all TE-derived reads with their corresponding annotations, and a structured matrix presenting TPM and read count data for each locus-specific TE. To mitigate the effects of TE homology, LATTE incorporated a pre-processing module for TE annotation filtration and a post-processing module for the detection of anomalous ERV expression. These modules are built upon long-read RNA-seq summaries and a machine learning-based outlier detection model, respectively (see Methods for computational details).

To allocate multi-mapped TE reads, we developed an assignment strategy based on an Expectation Maximization (EM) algorithm^33^ incorporating three indicators: subfamily identity, base coverage distribution, and genomic annotation region (**Figure 2C**). While existing tools often rely on a single indicator (e.g., gene expression) and exhibit moderate performance, LATTE leverages the principle that the distribution of multi-mapped reads is proportional to that of uniquely mapped reads across these specific indicators. This strategy expanded the probabilistic framework and optimized the iterative refinement process. In a typical scenario (**Figure 2C**), a multi-mapped read may align to three distinct regions belonging to different subfamilies. When the maximum probability exceeded a predefined threshold, the read was assigned to the corresponding genomic region, and the expectation step was updated for the remaining unassigned reads. This process continued iteratively until the threshold criteria were no longer met, at which point LATTE transitioned to a secondary indicator. This stage utilizes definite integration to compute initial probabilities based on the base coverage of each TE subfamily. If ambiguities persisted, a final indicator based on the TE annotation region was applied using the same computational approach. For reads that remain unassigned after evaluating all three indicators, the program initiated a new cycle to recalculate expectations within the pool of already assigned reads, terminating only when no further allocations could be made (see Methods).

### Benchmarking confirms superior accuracy and robustness of LATTE at locus-specific resolution

We benchmarked LATTE against three widely used tools based on EM or alignment (SQuIRE, TEtranscripts, and SalmonTE) and two baseline strategies: “Unique” (excluding all multi-mapped reads) and “Primary” (retaining only primary alignments, hereafter referred to as basic methods). Simulated RNA-seq data were generated from the human reference transcriptome (GCA_000001405.29) using wgsim^34^. Each simulated sample contains 20 million raw reads (including ∼6 million TE-derived reads) without base errors to eliminate biases stemming from data size or alignment quality. Additionally, we obtained ten real-world short-read RNA-seq data of human from the International Genome Sample Resource (IGSR)^35^ (**Supplementary Table1**) to evaluate tool stability by comparing the hierarchical structures^36^ of results derived from both simulated and real-world data.

First, we evaluated the computational performance of these tools using simulated data (**Figure 3A**). Execution times for EM-based tools were comparable, averaging approximately 60 minutes per sample (e.g., LATTE, SQuIRE, and TEtranscripts), while alignment-based or random assignment methods were significantly faster (e.g., SalmonTE at ∼16 minutes; basic methods at ∼5 minutes). Regarding disk I/O (“Block in” and “Block out”), LATTE provides three specific details for each TE read, genomic coordinates, TE subfamilies, and relative positions within the corresponding TE consensus, which are essential for its read-specific resolution. These details enable users to identify exact TE expression boundaries, which is critical for elucidating TE biology, such as the identifications of TE-derived transcripts^37^ and exon-TE junctions^38^.

**Figure 3.**
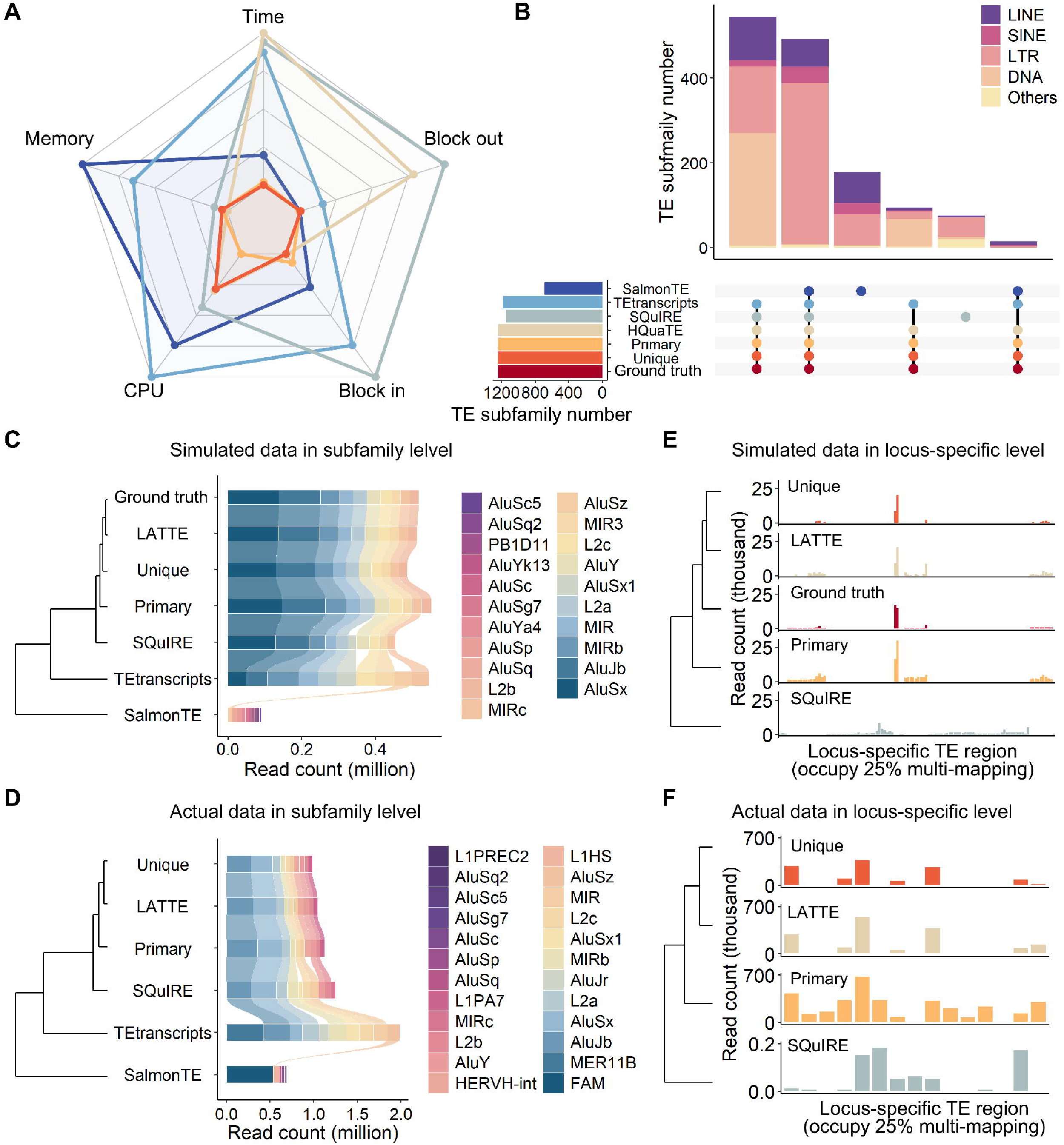
Benchmarking tools and strategies for TE quantification. (A) Computational resource consumption (CPU and memory). (B) UpSet plot showing detected TE types and their overlaps among tools and approaches. (C, D) TE quantification at the subfamily level for the top ten most abundant subfamilies in (C) simulated and (D) real-world data. (E, F) TE quantification at the locus-specific level, evaluated on regions harboring ∼25% of multi-mapped TE alignments in (E) simulated and (F) real-world data.

Next, we compared the handling of TE annotation files across tools (**Figure 3B**). TEtranscripts and SalmonTE rely on tool-specific annotation files, limiting their applicability primarily to model organisms like mouse, fruit fly, and zebrafish. In contrast, SQuIRE forced the collapsing of overlapping repeats in customized annotation files, which resulted in the omission of certain subfamilies (e.g., detecting only 1,148 subfamilies compared to 1,244 in the ground truth). LATTE and the basic methods directly utilized customized TE annotation files in GFF format, thereby preserving all TEs by default and allowing users to target specific subfamilies or regions. Consequently, this approach achieves the highest recall for subfamily capture (LATTE: 1.000; “Primary”: 1.000; “Unique”: 1.000) compared to others (TEtranscripts: 0.949; SQuIRE: 0.922; SalmonTE: 0.552), with 687 subfamilies (55.2%) consistently identified by all methods.

At the TE subfamily level, we analyzed the top ten most abundant subfamilies in simulated data (**Figure 3C**). Alignment-based quantification (SalmonTE) exhibited significant discrepancies in both composition and read counts, failing to align with the ground truth. Other tools also showed inconsistencies, such as L2b is accurately captured only by LATTE and the basic methods. Using the Intra-class Correlation Coefficient (ICC) to assess consistency with the ground truth, LATTE achieved the highest reliability (LATTE: 0.998; “Primary”: 0.983; “Unique”: 0.995; SQuIRE: 0.946; TEtranscripts: 0.198; SalmonTE: −0.256). These results were mirrored in real-world human IGSR data^35^ (**Figure 3D**), where alignment-based quantification showed a distinct bias in composition and read counts, likely due to mapping reads directly to TE consensus sequences. Notably, HERVH-int and L1PA7 were exclusively identified by LATTE and the basic methods. Tree-based clustering of read counts yielded consistent hierarchical structures across simulated and real-world datasets, validating the simulated data accurately captures real-world complexity and confirming the robustness of our evaluation framework.

Finally, we evaluated locus-specific quantification, a capability restricted to LATTE, SQuIRE, and the basic methods. We targeted genomic regions annotated by RepeatMasker^39^ containing homologous TE families—the primary source of multi-mapping ambiguity (**Supplementary Table2**). In simulated data (**Figure 3E**), 258 regions accounted for ∼25% of multi-mapped reads while covering less than 0.01% of the genome. Notably, SQuIRE tended to allocate reads to regions lacking uniquely mapped evidence, leading to nearly 50% incorrect assignments. Among the basic methods, “Unique” was overly conservative, while “Primary” was excessively inclusive. More specifically, while “Unique” performed best regarding ICC (0.883), and “Primary” achieved the highest recall (1.000), LATTE effectively balanced these extremes with an ICC of 0.839 and a recall of 0.944. In real-world data (**Figure 3F**), approximately 25% of the multi-mapped TE reads aligned to 15 specific regions. Likewise, LATTE maintained this balance, retaining eight fewer loci than “Primary” while capturing 31.4% more read counts than “Unique”. Tree-based clustering grouped LATTE and “Unique” together, separate from SQuIRE, further demonstrating LATTE’s accuracy in resolving locus-specific TE expression.

### Mitigating viral interference and sequence homology via machine learning and long-read integration

Endogenous retroviruses (ERVs) are a prominent subset of TEs^11^. In our previous study^5^, we observed a tenfold elevation in the expression of specific ERV types—notably RSV-int, RSV-LTR, and GGERVK10—in chicken samples. This finding is unexpected given that ERVs constitute only 1.84% of the chicken genome. We hypothesized that exogenous viruses might complicate TE quantification due to high sequence homology. To test this, we first confirmed significant sequence similarities (sequence length > 100 bp and BLAST identity > 90%) between ERVs and known exogenous viruses in chickens (**Figure 4A**). Notably, such high similarities were not observed in humans or other farmed animals (e.g., pigs, cattle, goats, and sheep). Furthermore, chicken samples infected with avian leukosis virus (ALV) exhibited significantly higher (*P* < 0.05) expression levels of EVAHP_I, RSV-int, and RSV-LTR compared to healthy control samples (**Figure 4B**; **Supplementary Table3**). These results confirm that exogenous viral sequences can lead to an overestimation of ERV expression through cross-mapping.

**Figure 4.**
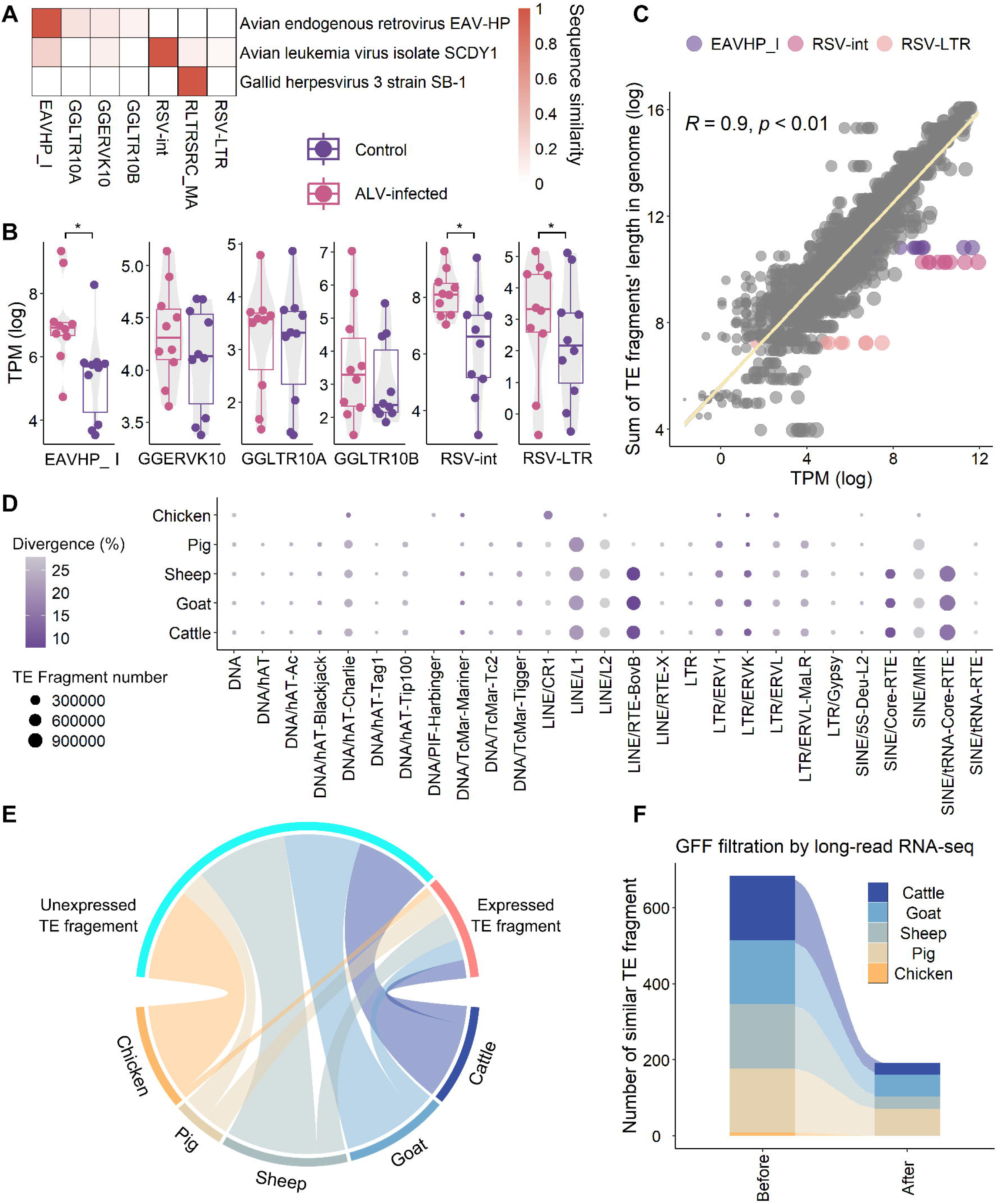
Mitigating sequence similarity effects with LATTE. (A) Heatmap illustrating sequence similarity between ERVs and exogenous viruses in chickens. (B) Differential expression of ERV subfamilies between ALV-infected and control samples, with each point represents an individual sample. Statistical significance was determined using an unpaired t-test with unequal variance (*P* < 0.05). (C) Linear regression analysis between expression level and genomic length, with anomalous ERVs highlighted as outliers. (D) Total number and divergence of TE fragments in common farm animal genomes, grouped by TE family. (E) Comparison of transcriptionally active versus silent TE subfamilies, revealing that only a small fraction of TEs is active. (F) Comparative analysis of genomic similarity before and after filtering transcriptionally silent subfamilies.

We observed a strong linear correlation (*R²* = 0.9, *P* < 0.01) between TE expression levels and their total genomic insertion lengths across most subfamilies. However, anomalous ERVs consistently appeared as outliers in this relationship (**Figure 4C**). To systematically identify these anomalies, we developed an optional module in LATTE using the Density-Based Spatial Clustering of Applications with Noise (DBSCAN) machine learning algorithm^40,41^. Briefly, this module calculates the ratio of expression level to total genomic insertion length for each subfamily. A DBSCAN model is then trained on these ratios from non-ERV families to establish a baseline for endogenous transcription. By applying this trained model to ERV families, the module can effectively predict and flag outliers likely caused by exogenous interference. This denoising module was validated using 16 chicken RNA-seq samples (eight ALV-infected and eight controls; **Supplementary Table3**), demonstrating its efficacy in isolating TE signals from viral noise.

Beyond viral interference, high sequence similarity and wide interspersion among TE families present additional challenges in their expression quantification (**Figure 4D**). This phenomenon, rooted in the intrinsic homology and transpositional mobility of TEs across diverse animal lineages^5^, often leads aligners to incorrectly map reads to transcriptionally silent regions. To address this, LATTE leverages long-read RNA-seq data, which unambiguously identifies expressed TE subfamilies by capturing full-length transcript sequence. We systematically analyzed 29 long-read RNA-seq datasets (**Supplementary Table4**) from five major livestock and poultry species, pigs (*n*=6), chickens (*n*=10), cattle (*n*=5), goats (*n*=4), and sheep (*n*=4), to catalog their transcriptionally active TE subfamilies. While 37.3 to 55.5% of genomic subfamilies exhibited activity, filtering TE annotation files against this active set refined the records, retaining only a high-confidence subset of expressed TEs (6.1 to 11.9% of original genomic records; **Figure 4E**). This refinement substantially reduced the number of TE fragments with high sequence similarity (**Figure 4F**), thereby facilitating the accurate assignment of multi-mapped reads. Currently, this long-read integration feature is optimized as an optional module specifically for domestic animals, aiming to maximize the detection efficacy and reliability of TE identification within the FarmGTEx project. We will consider more long-read RNA-seq datasets once they are available in the near future.

### Locus-specific TEs exhibit profound tissue-specificity and independent expression pattern

To assess the scalability of LATTE for quantifying locus-specific TE expression in domestic animals, we analyzed 250 short-read RNA-seq samples across five species (cattle, sheep, goats, pigs, and chickens; *n*=50 per species), encompassing ten distinct tissues (ovary, testis, spleen, skin, kidney, brain, lung, heart, liver, and muscle; with *n*=5 per tissue; **Supplementary Table5**). LATTE demonstrated remarkable efficiency, typically processing approximately one million TE-derived reads within ten minutes per CPU core (**Figure 5A**). Parallel gene expression quantification was performed using Stringtie^42^. Our analysis revealed striking tissue-specificity in locus-specific TE expression, with approximately 74.6% of TE loci expressed in only one tissue type (**Figure 5B**). Notably, the tissue-specificity of TEs significantly exceeded that of genes (**Figure 5C**). While 55.3% of genes were ubiquitously expressed across all ten tissues, only 13.3% of TEs showed such broad expression (**Figure 5C**). By overlapping TE coordinates with gene annotations (**Figure 5D**), we classified expressed TEs into “intragenic” or “intergenic” categories. We found that 94.3% of the locus-specific TE expressions were embedded in annotated gene regions (“intragenic”), while 5.7% were recognized as unannotated (“intergenic”). Intragenic TEs exhibited significantly lower sequence divergence (*P* < 0.01) compared to intergenic ones, suggesting they represent younger, more evolutionarily active insertions (**Figure 5E**). Furthermore, intergenic TE expression was predominantly captured in reproduction-related tissues, such as the ovary and testis (4.9%), compared to a minimal proportion in muscle (1.8%; **Figure 5F**).

**Figure 5.**
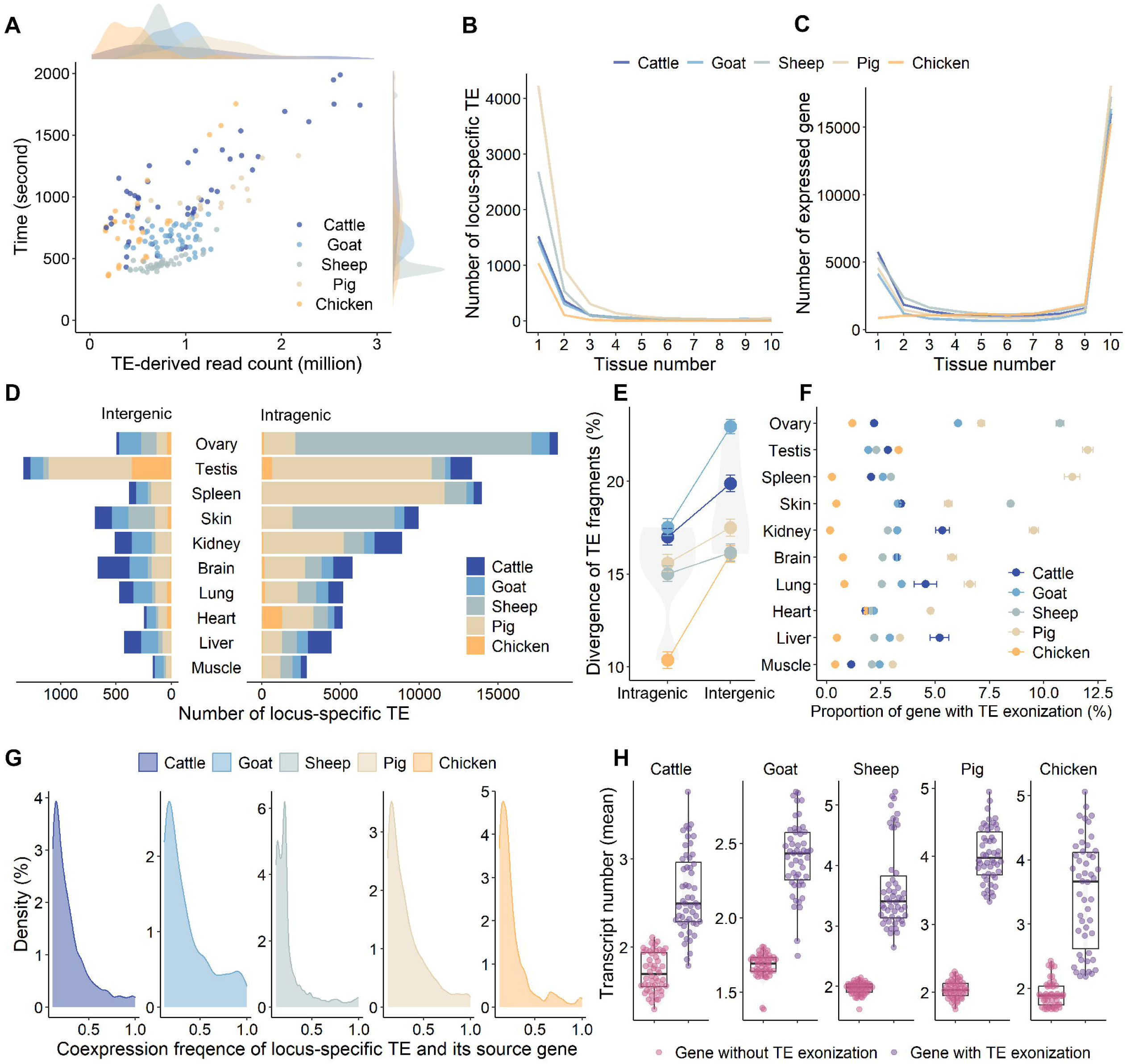
Characteristics of locus-specific TE expression. (A) Execution time of LATTE relative to the number of TE reads, processing approximately one million TE-derived reads per CPU core every ten minutes. Each node represents a sample, with color indicating the species. Tissue-specific expression patterns of (B) locus-specific TEs and (C) genes. (D, E) The total number of locus-specific TEs and their divergence patterns. TEs and host genes are linked based on genomic coordinate overlaps. TE divergence scores are provided by RepeatMasker^39^. (F) Proportion of genes exhibiting TE exonization. (G) Co-expression frequency of locus-specific TEs and their corresponding host genes across tissues. (H) Significant differences in transcript numbers between genes with and without TE exonization.

We further investigated the relationship between TEs and host genes by analyzing their co-expression patterns among tissue types, specifically focusing on TE exonization events. Density plots revealed that 38.7% of these exonization events occurred in only one tissue type, with a peak distribution at 0.1 (**Figure 5G**). These findings demonstrate that locus-specific TE exonization exhibited profound tissue specificity, even when the host gene itself is ubiquitously expressed. Given that TE exonization is functionally linked to tissue-specific alternative splicing^43^, we analyzed transcript diversity across the five studied species. We observed significant differences (*P* < 0.05) in the number of transcripts between genes with and without TE exonization (**Figure 5H**). Altogether, these findings highlight that the inclusion of TE sequences into mature transcripts is governed by highly localized regulatory programs rather than passive co-transcription.

### TE-eQTL reveal a genetic regulatory landscape decoupled from host genes

To evaluate the genetic regulatory mechanisms underlying TE expression, we collected RNA-seq data and corresponding genotype files from human lymphocyte (*n*=451), cattle blood (*n*=170), and chicken blood (*n*=192), and performed *cis*-TE-eQTL mapping following the PigGTEx project pipeline^18^ (**Figure 6A; Supplementary Table1**). First, expression values for locus-specific TEs were normalized using the trimmed mean of M-value (TMM) method, based on read counts and TPM values provided by LATTE. Second, we defined the *cis*-window for “intragenic” TEs as ±1_Mb around the transcription start site (TSS) of the host gene, and for “intergenic” TEs as ±1_Mb around their own genomic coordinates. Third, *cis*-TE-eQTL were mapped using OmiGA^44^, with nominal *P* adjusted via the Benjamini-Hochberg method^45^. Compared to previous strategies^15^, our pipeline offers two major advantages: (1) it utilizes each locus-specific TE as the basic quantification unit, enabling the discovery of differential genetic regulation within host genes; and (2) it supports *cis*-mapping resolution because LATTE provides precise genomic coordinates, thereby avoiding the high multiple-testing burden associated with genome-wide *trans*-scanning.

**Figure 6.**
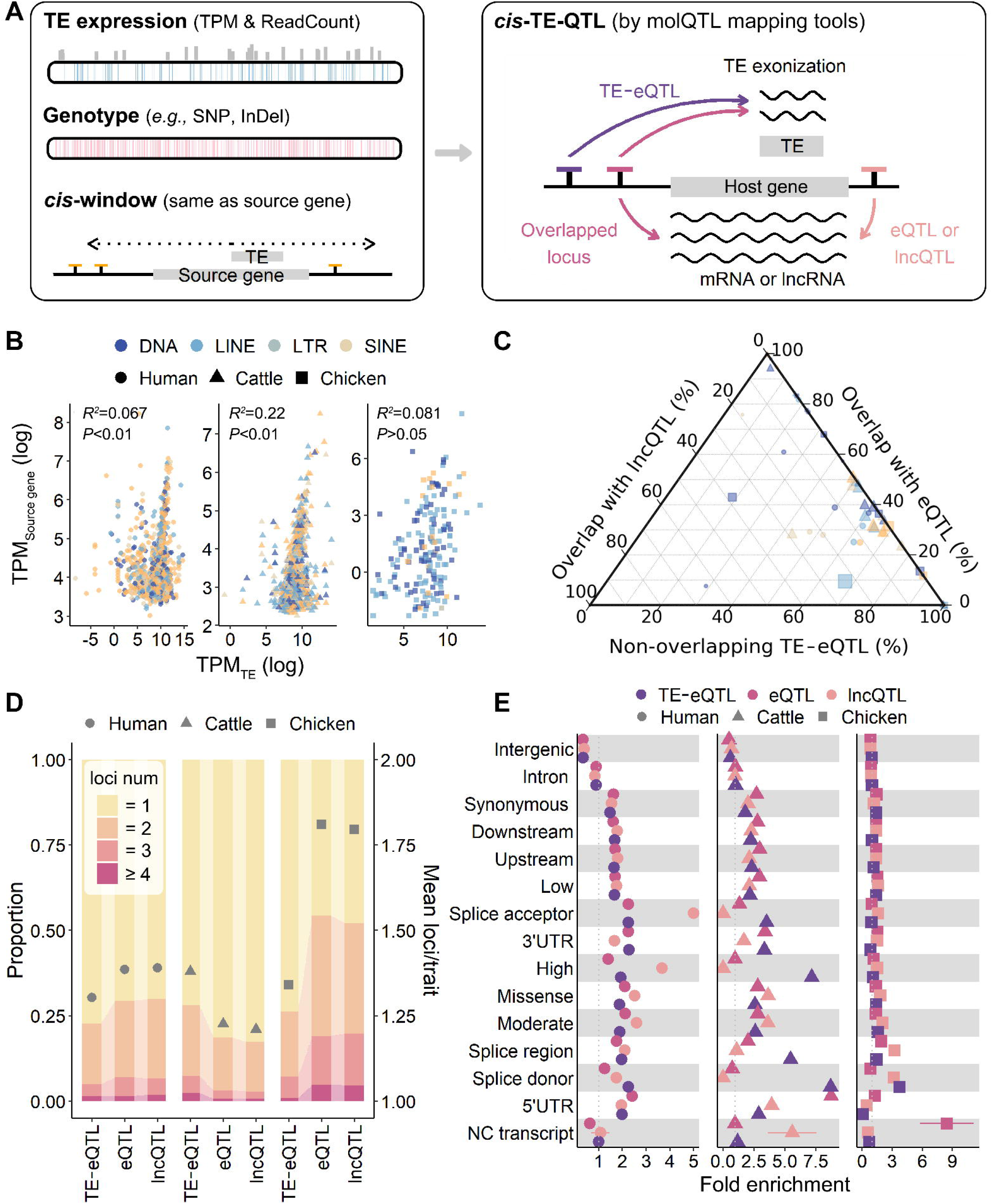
Discovery of genetic variants regulating TE expression. (A) Analytical framework for TE-eQTL mapping. The left diagram illustrates the identification of genetic variants associated with TE expression; the right diagram shows the overlap between TE-eQTLs, eQTLs, and lncQTLs. (B) Expression correlation between TEs and their host genes. Despite physical linkage via exonization, TE expression shows minimal correlation with host gene expression. (C) Proportion of TE-eQTLs overlapping with host gene QTL (eQTLs or lncQTLs). (D) Distribution of independent *cis*-QTL per molecular phenotype (TEs, PCGs, and lncRNAs) across species. (E) Functional annotation of TE-eQTLs, eQTLs, and lncQTLs using SnpEff^46^.

Analysis of genomic context revealed that 64.3%, 58.9%, and 57.8% of intragenic TEs were embedded within protein-coding genes (PCGs) in humans, cattle, and chickens, respectively, with a substantial portion also overlapping lncRNAs (25.1%, 8.6%, and 23.5%). To explore the relationship between host gene and TE expression, we quantified PCGs and lncRNA expression from the same datasets using Stringtie^42^ (**Supplementary Table1**). By pairing TEs with their host genes based on genomic coordinates overlaps, we observed notably weak expression correlations (*R²_human_*= 0.067; *R²_cattle_*= 0.22; *R²_chicken_*= 0.081; **Figure 6B**). Subsequently, we identified *cis*-eQTL and *cis*-lncQTL for these genes. Among significant loci (**Supplementary Table6**), only approximately 35% of *cis*-TE-eQTL were associated with PCGs at shared loci, and only 5% with lncRNA (**Figure 6C**). After accounting for linkage disequilibrium (LD), these proportions dropped to approximately 22% for PCG and 4% for lncRNA. These findings indicate that host gene expression did not accurately reflect the transcriptional activity of embedded TEs, pointing toward distinct, autonomous genetic regulatory mechanisms.

To further characterize this regulatory complexity, we identified independent signals for each gene and TE using conditional analysis in OmiGA^44^. The mean number of independent signals per PCG, lncRNA, and TE was 1.39, 1.39, and 1.28 in humans; 1.23, 1.21, and 1.38 in cattle; and 1.81, 1.79, and 1.34 in chickens (**Figure 6D**). To infer the potential functional impact of these signals (**Figure 6E**), we performed functional annotation using SnpEff^46^. Notably, *cis*-TE-eQTL enriched at splice donor sites across all studied species (enrichment factors—human: 2.3; cattle: 8.7; chicken: 3.7), suggesting a functional linkage between genetic variation, TE expression, and transcript splicing processes.

### TE-eQTL contribute to complex traits independently of host gene regulation

To systematically evaluate the relative contributions of TEs and genes to complex traits, we analyzed GWAS summary statistics from humans^47^, cattle^48^, and chickens^49^ (encompassing 133,441, 3,709, and 52,355 unique GWAS loci across 3,621, 55, and 70 traits, respectively). Using TWAS implemented in Fusion^50^, our analysis revealed that 200 human traits (5.5%) and 4 cattle traits (11.8%) were exclusively associated with TE expression (**Figure 7A**). For instance, the expression of AluJb and L2—both of which undergo exonization within the *DHFR* gene—was implicated in Huntington’s disease progression. This provided a novel mechanistic insight into a disease where *DHFR*’s role is well-documented but the involvement of its embedded TEs has been previously overlooked^51^. Overall, TE expression was associated with over half of the tested traits across all species (human: 53.3%; cattle: 64.7%; chicken: 53.3%; **Figure 7A**), underscoring its pervasive role in shaping phenotypic variation. Furthermore, we observed that vast majority of TWAS signals exhibited genomic independence (human: 94.6%; cattle: 72.7%; chicken: 90.2%), where TWAS-hit TEs and genes resided in distinct genomic regions (**Figure 7B**). Such evidence highlights TEs as an essential and unique component of the genetic architecture underlying complex traits.

**Figure 7.**
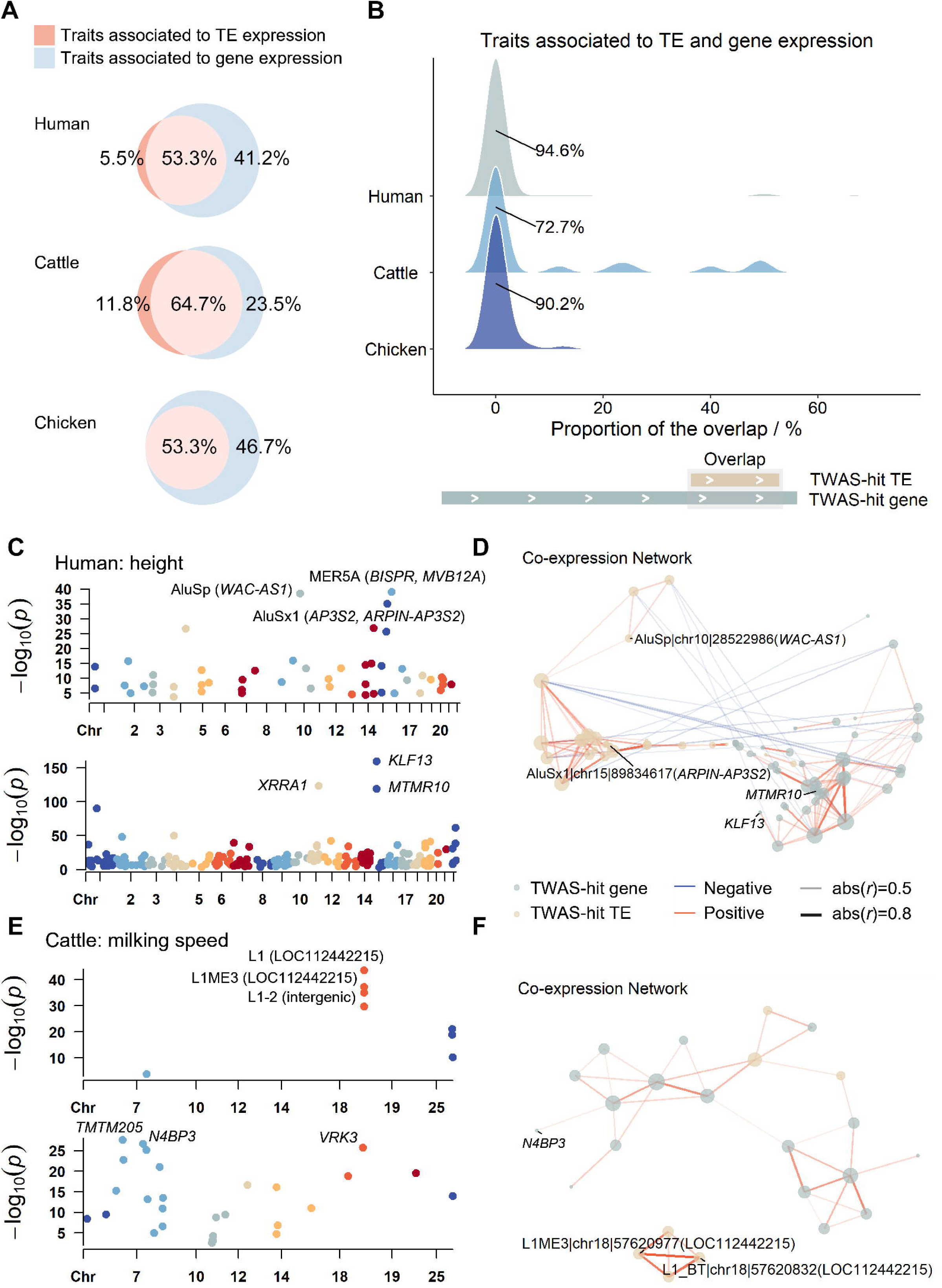
Interpreting complex traits with TE-eQTLs. (A) Proportion of complex traits associated with TEs, genes, or both in human, cattle, and chicken. (B) For traits associated with both TEs and genes, the proportion of TWAS loci where signals overlap genomically. (C, E) Manhattan plot of TWAS results for (C) human body height and (E) cattle milking speed, highlighting the top three associated TEs (host genes in parentheses) and genes. (D, F) Co-expression networks of TWAS signals for (D) human body height and (F) cattle milking speed (|r| > 0.4, *P* < 0.05). Node size represents degree of connectivity; edge width and color (red: positive; blue: negative) indicate correlation strength and direction.

To delineate the specific characteristics of these associations, we characterized the distinct genomic and expression patterns of the TWAS-hit TEs and genes. First, we analyzed their associations with human height (**Figure 7C**), where the top signals for TEs and genes exhibit high genetic independence. The leading TWAS-hit TEs included MER5A (overlapping *BISRP* and *MVB12A*), AluSp (overlapping *WAC-AS1*), and AluSx1 (overlapping *AP3S2* and *ARPIN-AP3S2*), while the top genes were *KLF13*, *XRPA1*, and *MTMR10*. A co-expression network analysis of these signals (|*r*| > 0.4, *P* < 0.05) revealed a clear bipartite structure (**Figure 7D**). This network separated into two discrete modules—one comprising co-expressed TWAS-hit TEs and another of co-expressed TWAS-hit genes—indicating independent transcriptional programs. Similarly, associations for cattle milking speed revealed distinct TWAS-hit TEs (e.g., L1, L1ME3, and intergenic L1-2) and genes (e.g., *TMEM205*, *N4BP3*, and *VRK3*; **Figure 7E**). These leading signals differed in both genomic coordinates and gene types, reinforcing the autonomous role of TEs. Furthermore, the corresponding co-expression network again split into two clusters based on feature type (**Figure 7F**). These results provide a novel perspective on the molecular regulatory processes, demonstrating that TEs represent a critical, independent layer of the functional genome.

### Splicing-mediated antagonistic regulation between TEs and host genes at the *IRF5* locus

Despite the predominant independence in genomic positioning and expression, we identified a subset of traits (human: 5.4%, cattle: 27.3%, chicken:9.8%) where TWAS signals from TEs and genes converged on shared genomic loci. Notably, at these overlapping loci, a portion of TE and gene associations exhibited opposite effect directions (human: 1.3%, cattle: 12.8%, chicken:4.2%), suggesting antagonistic regulatory relationships. A prominent example is the MSTB1-*IRF5* locus, where the TE and its host gene displayed opposite *Z* for Sjögren’s syndrome in humans (**Figure 8A**). This syndrome is a common manifestation of systemic lupus erythematosus (SLE)^52,53^. While previous studies established *IRF5* as a top risk gene for SLE^54–56^, and the human TWAS atlas confirmed its significant association across 20 tissues^57^, we discovered that the *cis*-TE-eQTL of MSTB1 exhibited stronger effect sizes compared to the *cis*-eQTL of *IRF5* (**Figure 8B**).

**Figure 8.**
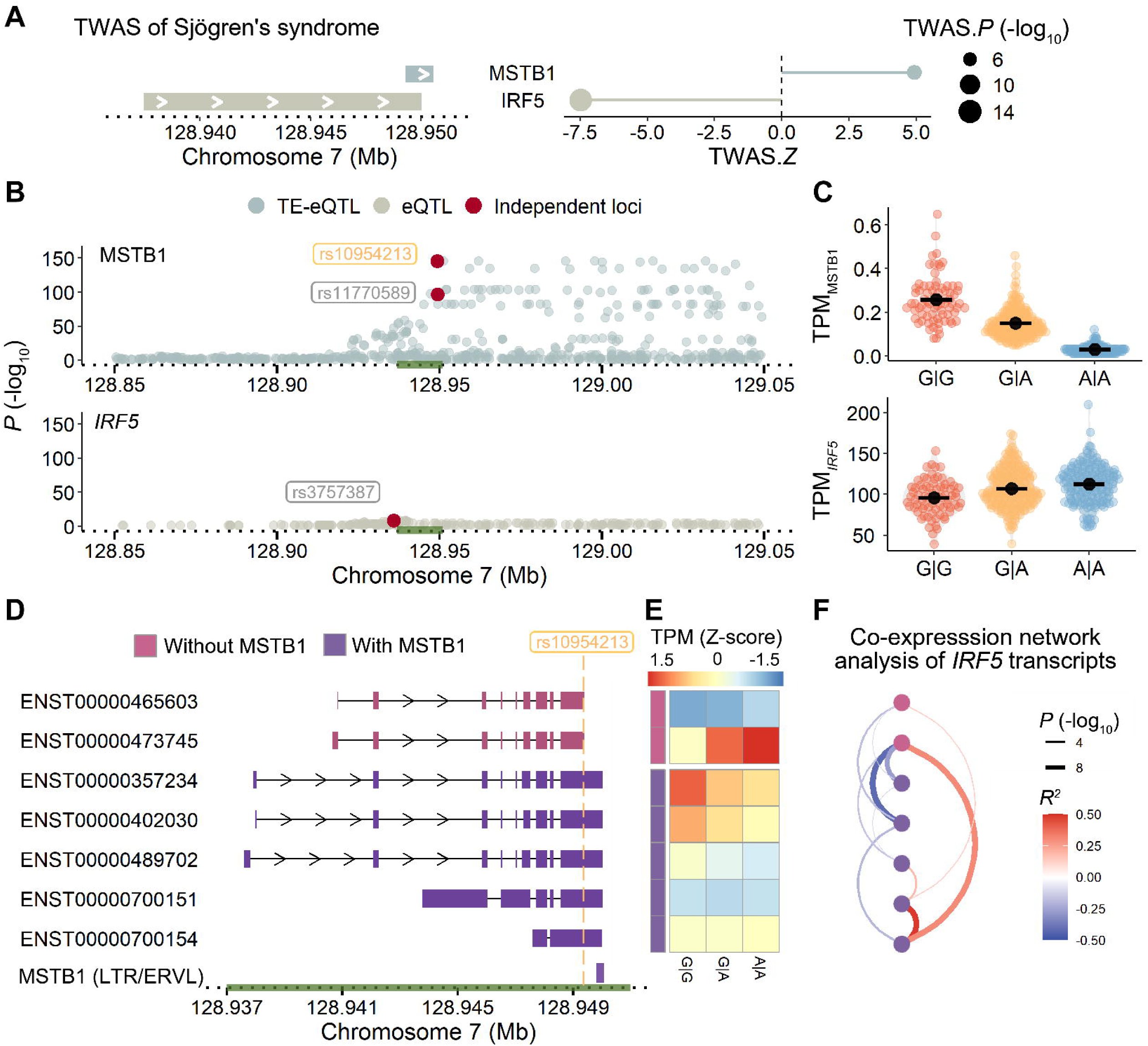
TE expression associates with complex traits via alternative splicing. (A) Genomic orientation of *IRF5* and MSTB1, and their TWAS associations with Sjögren’s syndrome. (B) *P* for eQTL (lower track, *IRF5*) and TE-eQTL (upper track, MSTB1). (C) Expression levels of MSTB1 and *IRF5*: boxplots demonstrate opposing regulatory directions of rs10954213 for the two phenotypes. (D) *IRF5* transcript architecture. (E) Heatmap of transcript expression levels across rs10954213 genotypes. (F) Co-expression network of the *IRF5* transcripts. Nodes are colored based on the inclusion of the MSTB1 sequence. Edge width and color represent the correlation *r^2^* and *P*, respectively.

We specifically focused on an independent *cis*-TE-eQTL of MSTB1 (rs10954213), which served as the lead SNP (cross-validation *R^2^*= 0.706) in our TWAS analysis. This mutation, located in the 3’UTR of *IRF5*, is a well-established genetic risk factor for SLE^58^. Our analysis revealed that rs10954213 exerts a modest but significant positive effect on *IRF5* expression while simultaneously negatively impacting MSTB1 (**Figure 8C**). To investigate this phenomenon, we examined the expression of individual *IRF5* transcripts. Among the seven *IRF5* transcripts identified, two contains MSTB1 exonization (**Figure 8D**). Heatmap analysis revealed that the rs10954213 risk allele up-regulates transcripts lacking MSTB1 while down-regulating those containing it (**Figure 8E**). Furthermore, a co-expression network confirmed that this antagonistic relationship is driven by the competition between the most abundant MSTB1-containing transcript (ENST00000473745) and the abundant transcripts lacking it (ENST00000357234 and ENST00000402030; **Figure 8F**). These findings demonstrated that the antagonistic relationship between TEs and host genes can be mediated by alternative splicing and transcript abundance shifts.

## Discussion

LATTE bridges a critical gap in functional genomics by moving beyond the simple annotation of TE sequences to directly quantify their transcriptional activity and phenotypic impact. It addresses the field’s primary obstacle—TE mobility and sequence homology—through an innovative reassignment strategy powered by an EM algorithm and denoising solutions, integrated with long-read sequencing and machine learning techniques. By providing read-specific resolution, this tool empowers researchers to uncover the patterns of locus-specific TEs and delineate the complex, often independent, relationships between TEs and host genes. Our findings demonstrate that LATTE enables the precise quantification and localization of TE expression, serving as a robust engine for future biological discovery.

Despite the widespread study of TEs, computational challenges in their quantification persist. Most existing tools lack read-specific resolution—a crucial limitation for precise locus-specific quantification that prevents the definitive identification of host genes, exon-TE junctions, and TE-derived transcripts^38^. Additionally, many tools impose rigid requirements for TE annotation files^24–27^, limiting their utility across diverse species or for comprehensive TE analysis. LATTE overcomes these limitations through a key innovation in reassigning multi-mapped reads. While current EM-based tools^25–27^ often rely on a single indicator and exhibit biases in simulations, LATTE implements a tiered, multi-indicator reassignment strategy. By iteratively applying three distinct indicators to assign reads, LATTE achieves a more than threefold improvement in reassignment efficacy compared to single-indicator methods.

The inherent homology of TEs remains a primary obstacle to accurate alignment. Our study confirms that significant sequence similarity exists not only among dispersed TE copies but also between ERVs and exogenous viruses, leading to a high proportion of multi-mapped reads^59^. LATTE addresses these challenges by employing a DBSCAN-based machine learning module^40,41^ to identify and flag anomalous ERV expression likely caused by viral interference. Furthermore, we leverage long-read RNA-seq data to catalog transcriptionally active TE subfamilies, creating a high-confidence filtering set for TE annotations. These embedded modules minimize the biases introduced by TE homology, ensuring that the quantified signals reflect true endogenous transcription. Currently, this long-read integration feature has been fully optimized for major domestic animals, providing comprehensive annotation resources to support high-resolution TE quantification studies in these species.

A particularly intriguing finding of our study is the subset of traits (human: 1.3%, cattle: 12.8%, chicken: 4.2%) where TWAS signals for TEs and host genes converge on shared loci but exert antagonistic effects. As exemplified by the MSTB1*-IRF5* locus, this opposing regulation is mediated by splicing-induced transcript competition. While recent studies have highlighted the role of exon-TE junctions in tumor biology^38^ and protein isoforms diversity^22^, they often focused on transcripts with TE insertions, overlooking the independent genetic regulation of TEs versus host genes^60^. This antagonistic relationship highlighted a sophisticated layer of genomic regulation, where TEs are not merely passive drivers of variation but active participants in balancing regulatory networks, potentially serving as a fine-tuning mechanism for complex traits and diseases.

Despite these advancements, LATTE has certain limitations. First, its accuracy in assigning multi-mapped reads is partially contingent upon the presence of uniquely mapped reads at a given locus; regions entirely lacking such unique signals remain challenging. Second, the precision of read-specific resolution can be sensitive to RNA-seq datasets with high mismatch rates. Furthermore, while our study identifies traits exclusively linked to TE expression, the observational nature of the data precludes definitive causal inferences without further experimental validation.

In conclusion, LATTE demonstrates exceptional precision in identifying the genomic origins of individual TE reads, enabling sophisticated analyses such as *cis*-TE-eQTL mapping. As a foundational component of the international FarmGTEx project, LATTE has established a standardized pipeline for repetitive element analysis. Its scalability and cross-species accuracy position it as an essential resource for future large-scale functional genomic efforts, aiming to unlock the “dark matter” within the genetic architecture of domesticated and diverse species, and bridging the knowledge gap between TE expression and complex phenotypic variation.

## Materials and Methods

### Software development and implementation

To ensure LATTE is as user-friendly as possible, we streamlined the input and output requirements. Users only need to provide two input files: a coordinate-sorted BAM file from RNA-seq data and a TE annotation file (GFF format) specifying the genomic coordinates of TEs. The tool supports both paired-end and single-end RNA-seq reads. For TE annotation, we recommend using RepeatMasker (v4.1.6)^39^ to identify TEs within the reference genome. Alternatively, users may provide custom GFF files, provided they contain the following essential tab-delimited columns: (1) chr (Column 1): chromosome name (e.g., chr1, chrX); (2) start (Column 4): starting genomic position of the TE insertion; (3) end (Column 5): ending genomic position of the TE insertion; and (4) TE info (Column 9): identity of the TE, typically including the TE subfamily. Non-essential columns can be filled with placeholder values to maintain the standard format. Detailed input specifications and examples are available on our GitHub repository (https://github.com/PengjuZ/LATTE).

The LATTE pipeline consists of three core stages: (1) **Preprocessing**, which includes classification of read types, recognition of TE reads, and filtration of TE annotation file; (2) **Localization**, which assigns the multi-mapped TE reads to high-confidence genomic positions; and (3) **Post-processing**, including the detection of anomalous ERV expression. Among them, the filtration of TE annotation file and the detection of anomalous ERV expression are optional modules. By default, LATTE generates two primary output files, redirected to the working directory if no output parameter is provided. The first output file, suffixed with *_TE file, includes the TE read’s coordinate and its overlapped TE annotation. The second output file, suffixed with *_summary, contains a statistical matrix with three columns: TE annotation, TPM, and read count. Notably, the row count and order of this file are identical to those in the provided GFF file, enabling users to perform straightforward column-wise concatenation for TE-eQTL mapping.

### Classification of read types

LATTE categorizes read types based on information encoded in standard BAM files. First, alignment tags (typically found in the 12th column of a BAM file) are used to distinguish uniquely mapped reads from multi-mapped reads. Specifically, the tool utilizes “NH:i:N” tag, which indicates the total number of reported mapping locations for a read or read pair. Uniquely mapped reads are defined by the “NH:i:1” tag, signifying alignment to a single genomic position. Using SAMtools (v1.17)^29^, LATTE executes the filter expressions [AS] && [NH]==1 and [AS] && [NH]!=1 via the samtools view command to partition uniquely mapped and multi-mapped reads, respectively. Second, LATTE adopts individual segments of splice junction reads as its fundamental quantitative unit (**Figure S1A**). To achieve this, the tool parses the Compact Idiosyncratic Gapped Alignment Report (CIGAR string, the 6th column of a BAM file) to distinguish between spliced junction reads and full-length reads. For example, a 150 bp read that is continuously aligned is characterized by a CIGAR string of “150M”. In contrast, a read spanning two exons separated by a 1000 bp intron is represented as “75M1000N75M”. For subsequent quantification, LATTE employs the bedtools bamtobed-split command to decompose these spliced junction reads into discrete segmental records, while treating full-length reads as individual records.

### Recognition of TE reads

LATTE identifies TE-derived reads by overlapping their alignment coordinates with the annotated TE loci. However, due to the fragmented nature of TEs within the genome (**Figure S1B**), the minimum overlap requirement between the TE annotation file and reads’ genomic coordinate file significantly affects TE reads quantification^59^. This often leads to substantial inconsistencies between studies utilizing different thresholds—specifically, more stringent overlap requirements typically result in lower TE read counts. By default, LATTE applies a 0.5 overlap threshold, meaning a read is only assigned to a TE if at least 50% of its length overlaps with a single TE annotation (**Figure S1C**). Users can increase the sensitivity of TE detection by lowering this threshold, which allows for a higher percentage of the read to map to flanking sequences (**Figure S1D**). This flexibility provides a granular view of TE fragments and is particularly advantageous for defining the precise boundaries of TE exonization, such as the junctions between TEs and neighboring exons.

### Localization of TE reads

Following the preprocessing steps, TE-derived reads are categorized into two distinct groups: multi-mapped and uniquely mapped. Here, uniquely mapped reads serve as the foundational data for estimating the relative expression across three hierarchical indicators: TE subfamilies, base coverages within consensus sequences of each subfamily, and individual genomic annotations. The efficacy of the EM (Expectation-Maximization) assignment algorithm relies on the assumption of relative consistency in these indicators between uniquely mapped and multi-mapped reads. Notably, these indicators are arranged in a hierarchical order, progressing from broader genomic regions to specific loci. The computational processes for these indicators are respectively elucidated below.

First, TE subfamily-level assignment. Assuming a sample contains several TE subfamilies (*i*). LATTE starts by aggregating the read counts for each TE subfamily from uniquely mapped reads (*Readcount_i_^single^*). By incorporating the total read count of multi-mapped TE reads (*Readcount_total_^nultiple^*), the expected read count of each subfamily (*Expectation_i_*) is calculated. Subsequently, LATTE calculates the initial probability distribution across TE subfamilies (*P_i_^initial^*): defined as the ratio between expected read count and the actual multi-mapped read count. Finally, through normalization by the potential subfamilies for a given multi-mapped TE read, these normalized values are utilized to assign reads that satisfy the minimum threshold. Reads falling short of this threshold undergo further iterations by other indicators.

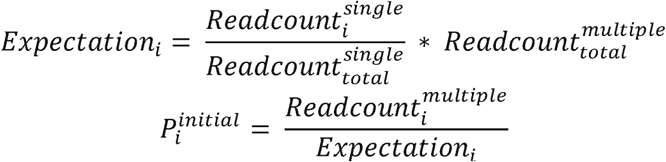

Second, consensus sequence-level assignment. LATTE utilizes the relative positioning of TEs within the consensus sequence of each TE subfamily. This step posits that sections of the TE consensus with higher coverage and depth, as established by uniquely mapped reads, have a higher probability of assignment. The process begins by calculating the TE subfamily types (i) and their respective consensus coordinates. It then uses an overlapping technique to map genomic coordinates of reads to relative positions within the TE consensus. The uniquely mapped reads are used to determine the coverage and depth of each TE consensus. For multi-mapped reads, LATTE identifies their overlapping start (*start*) and end (*end*) positions within the TE consensus, along with their corresponding depth (*BaseDepth_i_^nultiple^*). The initial probabilities (*P_i_^intial^*) for a given multi-mapped TE read at different loci are then computed using a definite integral format. In this step, reads that fail to meet the minimum threshold requirement undergo further iterations as well.

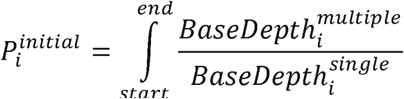

Third, annotation-level assignment. LATTE refines the genomic range for probability calculation by utilizing each TE annotated region as a fundamental unit of analysis. The assignment of multi-mapped TE reads is based on the specific read counts of individual annotations, as determined by uniquely mapped reads. This step follows the same core logic as the first indicator but operates at the resolution of specific genomic loci. In cases where a multi-mapped read still fails to meet the threshold, it is subsequently re-processed by the first indicator to ensure maximal signal recovery.

### Filtration of the TE annotation file

This optional module, integrated into the filterGFF script, is designed to mitigate the biases introduced by sequence homology among genomic TEs. The module functions by cross-referencing TE subfamily identifiers between a user-provided text file of transcriptionally active subfamilies and the master TE annotation file. Notably, while we have validated this functionality primarily using GFF-formatted files from RepeatMasker (v4.1.6)^39^, users may adapt other custom files to this format.

The list of active TE subfamilies was derived from long-read RNA-seq data (**Supplementary Table4**), which was selected for its superior capacity to capture full-length TE transcripts with high confidence. This approach establishes a robust baseline for confirming the transcriptional activity of specific TE subfamilies. Currently, pre-compiled active subfamily lists are available for major livestock or poultry species, including pigs, cattle, sheep, goats, and chickens.

### Detection of anomalous ERVs expression

This optional module is designed to efficiently identify potential interference induced by exogenous viruses. The process consists of three main stages: (1) Extracting TE expression at the subfamily level (n) and aggregate genomic length for each subfamily based on the TE annotation file; (2) Calculating the ratio between total expression and aggregate length for each TE subfamily; and (3) Applying the DBSCAN algorithm from Scikit-learn (v1.5.1)^32^ to identify statistical outliers. Finally, samples exhibiting outlier ratios are classified as having anomalous ERV expression and are recorded in a separate output file. The algorithm utilizes Euclidean distance as the primary metric with a default epsilon (eps) value of 3. Users may adjust these parameters within the Python script to fine-tune detection sensitivity according to their specific requirements.

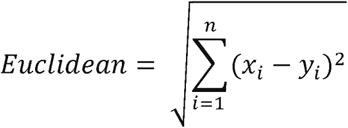

### TE annotation and sequence analysis

The reference genome assemblies utilized in this study included human (GCA_000001405.29), pig (GCA_000003025.6), chicken (GCA_016699485.1), cattle (GCA_002263795.3), sheep (GCA_016772045.1), and goat (GCA_016772045.1). TEs within these reference genome were identified using RepeatMasker (v4.1.6)^39^ with the resulting GFF files selected as the primary TE annotation source. Notably, we recommend filtering out low-complexity sequences, as their retention can substantially impair computational efficiency.

TE sequences were retrieved from Repbase (v20170127)^61^ and subsequently formatted into FASTA format files for downstream analysis. Host-specific viral sequences were acquired from EBI (https://www.ebi.ac.uk/genomes/virus.html). We employed BLAST (v2.5.0)^62^ to conduct sequence similarity searches by species. The process began with the construction of a nucleotide database using the FASTA file containing all TE sequences. Subsequently, BLASTN (v2.5.0)^62^ was utilized to compare the sequences, generating a series of output files. These outputs were systematically organized by TE subfamilies specific to each species. To ensure high-quality results, we reported only those sequences with an identity exceeding 90% and an alignment length greater than 100 bp. TE divergence scores for each TE fragment were extracted from the RepeatMasker (v4.1.6)^39^ output, where higher divergence levels served as a proxy for greater evolutionary age. Finally, locus-specific TE expression was classified as either “intragenic” or “intergenic” based on the degree of overlap with reference gene models.

### Publicly available data acquisition

The benchmarking of different methods was conducted using human RNA-seq samples (**Supplementary Table1**). We further validated the method’s accuracy and scalability using a dataset of 250 RNA-seq samples derived from five major farm animals (cattle, sheep, goats, pigs, and chickens), with 50 samples per species (**Supplementary Table5**). This comprehensive dataset spans ten distinct tissues—lung, liver, heart, ovary, brain, muscle, testis, spleen, skin, and kidney—with five replicates per tissue. Additionally, to evaluate the detection of anomalous ERV expression, we analyzed data from eight ALV-infected and eight control chickens (**Supplementary Table3**).

TE-eQTL and eQTL mapping were performed on human lymphocyte cells (*n*=451), cattle blood (*n*=170), and chicken blood (*n*=192). Specifically, human samples were retrieved from the International Genome Sample Resource (IGSR)^35^, while cattle and chicken samples were obtained from our previously published studies. The metadata for all samples is provided in **Supplementary Table1**.

The GWAS summary statistics are detailed as follows. (1) Human^47^: A resource built by integrating genetics from the GWAS Catalog and UK Biobank with multi-omics data (transcriptomic, proteomic, and epigenomic data). This comprehensive database encompasses 133,441 published GWAS loci across 3,621 complex traits and diseases. (2) Cattle^48^: Analysis based on 294,079 first-lactation Holstein cows, covering 3,709 loci associated with 55 traits, including production, fertility, and somatic cell score. (3) Chicken^49^: Data derived from the chicken GTEx project, comprising 52,355 loci linked to 39 complex traits, including growth, carcass quality, egg production, feed efficiency, and blood biochemical indices.

### Short-read RNA-seq data processing

To ensure a robust evaluation, we conducted a benchmarking analysis utilizing both real-world and simulated data. Real-world RNA-seq dataset were sourced from the EBI SRS server^63^ and GEO database^64^. Raw data were downloaded and converted to FASTQ format using the SRA toolkit (v3.1.1)^65^. Quality control was performed using FASTQC (v0.12.1)^66^ to ensure the removal of low-quality reads. The resulting high-quality reads were then aligned to the reference genome using STAR (v2.7.11a)^67^.

For the simulation-based benchmark, we generated ten independent datasets of 150 bp paired-end RNA-seq reads from the human reference transcriptome (GCA_000001405.29) using wgsim (v1.0)^34^. Each dataset comprised 20 million read pairs to provide sufficient depth for the EM algorithm’s estimation. The base error rate was set to zero to eliminate sequencing errors as a source of alignment ambiguity, thereby isolating alignment ambiguity as the primary variable for benchmarking. The ground-truth genomic coordinates were extracted from the read headers generated by wgsim. Subsequent alignment of simulated data was also performed with STAR (v2.7.11a)^67^.

For QTL mapping, we then extracted the raw read counts of genes by featureCounts (v2.0.6)^68^ and obtained their normalized expression (TPM) using Stringtie (v2.2.1)^42^. The identification of splice junctions between exons and TEs followed previously established protocols^22^. The expression of transcript is identified by its TPM and coverage using Stringtie (v2.2.1)^42^, restricted to the provided reference gene and transcript annotations.

### Benchmarking of TE quantification

As described in the introduction, diverse strategies have been developed to estimate TE expression. In this study, we evaluated three prominent tools capable of quantifying all TE classes: SalmonTE^24^, TEtranscripts^26^, and SQuIRE^25^. Furthermore, two baseline quantification strategies were included for comparison: one that exclusively considers uniquely mapped TE reads (“Unique”), and another that assigns multi-mapped reads to their primary alignments (“Primary”).

To assess the computational performance of these approaches and tools^24–26^, we systematically monitored execution time, peak memory usage, CPU utilization, and disk I/O. These metrics provide a comprehensive framework to quantify processing efficiency and identify performance bottlenecks, facilitating an objective assessment of each tool or method.

To evaluate performance at the subfamily level, we utilized the quantified read counts for each TE subfamily as the primary metric, from which mean and standard deviation of expression levels were calculated. For locus-specific benchmarking, we compared read counts within specific genomic regions (**Supplementary Table2**). Notably, these regions accounted for approximately 25% of all multi-mapped TE alignments, despite representing less than 0.01% of the total genome.

For simulated data, the ICC was conducted to evaluate the concordance between ground truth values and tool estimates using the “irr” package (v0.84.1) in R (v4.3.1)^69^. Recall in our study was defined as the ratio of correctly captured signals to the total ground-truth count for TE subfamilies and locus-specific regions, respectively. Finally, to quantify the dissimilarity in read count estimates produced by different tools or methods, a pairwise Euclidean distance matrix was computed for both simulated and real-world datasets using the dist function in R (v4.3.1)^69^.

### Identification and annotation of *cis*-molecular QTL

For molecular QTL (molQTL) mapping, we utilized human lymphocyte cells (*n*=451), cattle blood (*n*=170), and chicken blood (*n*=192). We considered variants with Minor Allele Frequency (MAF)□≥□1% and minor allele count□≥□6, resulting in 73.5 million, 12.1 million, and 14.5 million variants for humans, cattle, and chickens, respectively.

To control for population effects and technical noise on the discovery of QTL^70^, we utilized five genotype Principal Components (PCs) and ten gene expression PCs as covariates to account for hidden confounders. The TE-eQTL and eQTL mapping procedures were performed following the protocols established by the PigGTEx project^18^. Expression levels of protein-coding genes (PCGs), lncRNA genes, and locus-specific were normalized using the Trimmed Mean of M-value (TMM) method in edgeR (v3.14.0)^71^. Genes or TEs with TPM < 0.1 and a read count < 6 in more than 80% of samples were excluded from the analysis.

We defined the *cis*-window of a gene as ±1_Mb around its transcription start site (TSS). For “intragenic” locus-specific TEs, the *cis*-window was defined as the ±1_Mb region around the TSS of the host gene, ensuring a consistent genomic context for subsequent integrative analyses. For “intergenic” locus-specific TEs, we used the ±1_Mb window relative to their own genomic start sites as recorded in the TE annotation file. Significant associations were identified using the *cis*-mode of OmiGA (v0.5.3.241214_beta)^44^. we employed the *cis* mode to identify significant associations of variants with gene expression. We used a permutation approach to identify genome-wide significant genes (eGenes) with a false discovery rate (FDR) of 0.05, determined by the Benjamini–Hochberg method^45^. For significant *cis*-molQTLs, a conditional analysis was performed using the independent-*cis* mode in OmiGA to identify independent signals. Finally, independent *cis*-loci were annotated using SnpEff (v4.3.1t)^46^ to predict their potential biological effects against the human (GCA_000001405.29), cattle (GCA_002263795.3), and chicken (GCA_016699485.1) reference genomes.

### Transcriptome-wide association studies of complex traits

To investigate whether the overall *cis*-genetic component of molecular phenotypes (including both genes and TEs) is associated with complex traits, we conducted transcriptome-wide association studies (TWAS) using Fusion^50^. We trained a predictive model for each TE-eGene (eTE) or eGene. To ensure model reliability, we only retained predictive models with a cross-validated value > 0.1 and a prediction performance *P* < 0.05 for further TWAS analysis. Finally, to assess each molecular phenotype’s association with traits, we performed TWAS by integrating these predictive models using GWAS summary statistics data from human^47^, cattle^48^, and chicken^49^ populations, as detailed in the preceding sections.

### Statistical analysis and visualization

All statistical analyses, including the calculation of means, standard deviations, and correlation coefficients, were performed using R (v4.3.1)^69^. Statistical significance was determined using appropriate tests for each dataset. Data visualization was primarily conducted using several robust R packages, including pheatmap (v1.0.12)^72^ for heatmaps, CMplot (v4.5.1)^73^ for Manhattan and QQ plots, patchwork (v1.3.0) for multi-panel figure assembly, and ggplot2 (v3.5.1)^74^ for general plotting. These tools ensured both the statistical rigor and the clarity of our graphical representations.

## Data and code availability

LATTE is an open-source tool freely available through GitHub (https://github.com/PengjuZ/LATTE), which includes the source code, detailed documentation, and example datasets. The source of the raw sequencing data in this study is recorded in supplemental tables. Additional scripts used for data integration and figure generation are also hosted in the GitHub repository to ensure the reproducibility of our findings.

## Author contributions

P.Z. and J.H. conceived and designed the experiments. J.H. implemented the LATTE software. P.Z., J.H. and C.P. designed an analytical strategy and performed analyses. Y.Z., H.Z. and Z.W. collected and prepared for sequencing data. J.H. wrote the initial version of the manuscript and P.Z., and L.F. contributed to the subsequent versions. All authors reviewed and approved the final version of the manuscript.

## Acknowledgments

This work is financially supported by the National Natural Science Foundation of China (Grant No. 32202626 and 32370678).

---

**Supplementary Figure 1.**
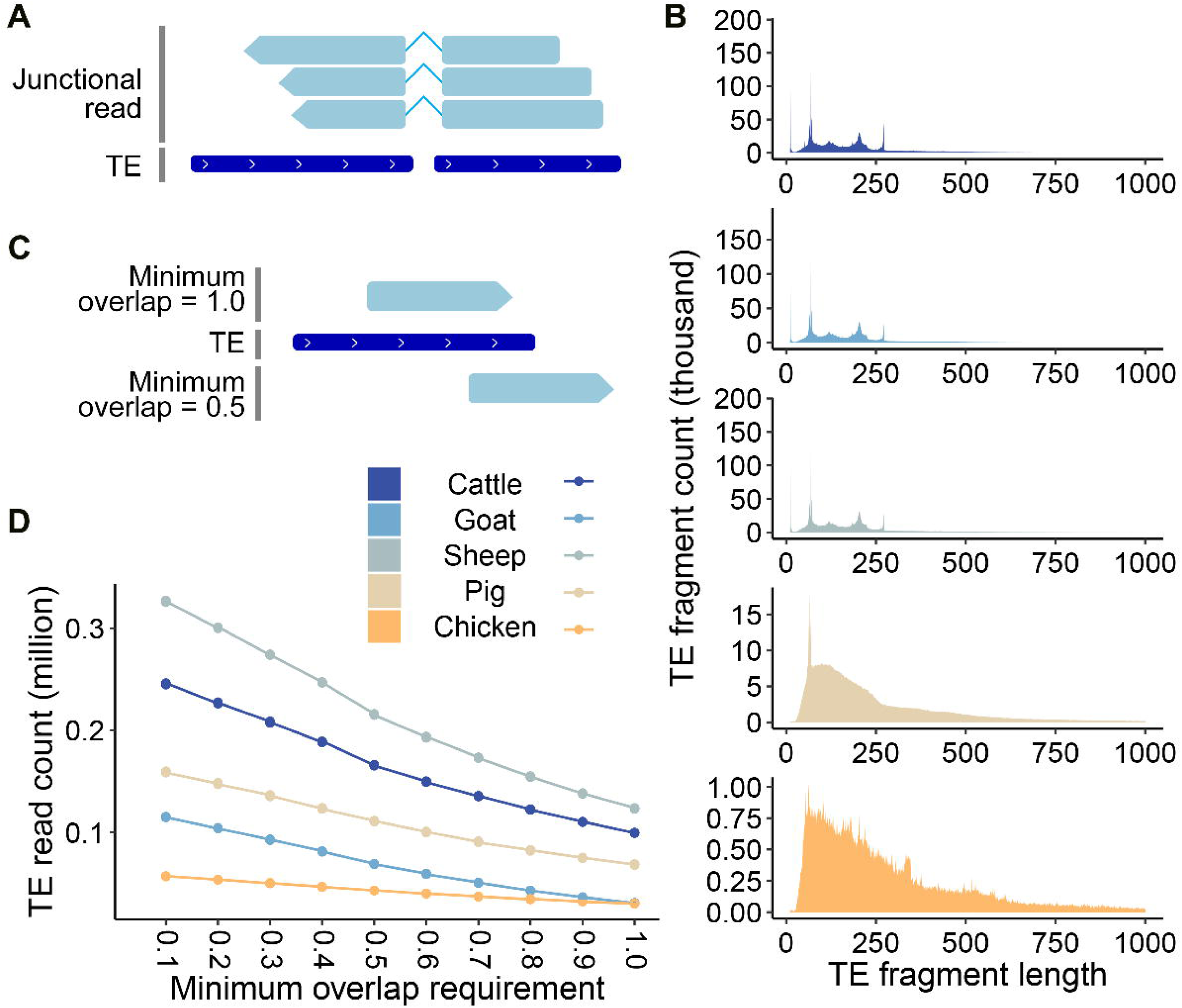
TE fragments in genome. (A) Schematic of the junctional reads spanning a splice site between a TE and a host gene exon. (B) Genomic length distribution of annotated TE fragments. (C) Illustration of the minimum overlap parameter, showing examples of a read fully within a TE annotation and a read with partial (50%) overlap. (D) The impact of the minimum overlap threshold on the number of reads classified as TE-derived.

## Reference

1. Fedoroff, N. V. Transposable Elements, Epigenetics, and Genome Evolution. Science 338, 758–767 (2012).

2. Wicker, T. et al. A unified classification system for eukaryotic transposable elements. Nat Rev Genet 8, 973–982 (2007).

3. Bourque, G. et al. Ten things you should know about transposable elements. Genome Biol 19, 199 (2018).

4. Cordaux, R. & Batzer, M. A. The impact of retrotransposons on human genome evolution. Nat Rev Genet 10, 691–703 (2009).

5. Zhao, P., Peng, C., Fang, L., Wang, Z. & Liu, G. E. Taming transposable elements in livestock and poultry: a review of their roles and applications. Genet Sel Evol 55, 50 (2023).

6. Du, A. Y., Chobirko, J. D., Zhuo, X., Feschotte, C. & Wang, T. Regulatory transposable elements in the encyclopedia of DNA elements. Nat Commun 15, 7594 (2024).

7. Hormozdiari, F. et al. Functional disease architectures reveal unique biological role of transposable elements. Nat Commun 10, 4054 (2019).

8. Lanciano, S. & Cristofari, G. Measuring and interpreting transposable element expression. Nat Rev Genet 21, 721–736 (2020).

9. Payer, L. M. & Burns, K. H. Transposable elements in human genetic disease. Nat Rev Genet 20, 760–772 (2019).

10. Rivas-Carrillo, S. D., Pettersson, M. E., Rubin, C.-J. & Jern, P. Whole-genome comparison of endogenous retrovirus segregation across wild and domestic host species populations. Proc. Natl. Acad. Sci. U.S.A. 115, 11012–11017 (2018).

11. Tokuyama, M. et al. ERVmap analysis reveals genome-wide transcription of human endogenous retroviruses. Proc. Natl. Acad. Sci. U.S.A. 115, 12565–12572 (2018).

12. Lei, X. et al. ERVcancer: a web resource designed for querying activation of human endogenous retroviruses across major cancer types. Journal of Genetics and Genomics S1673852724002418 (2024) doi:10.1016/j.jgg.2024.09.004.

13. Yu, J. et al. Endogenous retrovirus activation: potential for immunology and clinical applications. National Science Review 11, nwae034 (2024).

14. Zhou, B. et al. Endogenous Retrovirus-Derived Long Noncoding RNA Enhances Innate Immune Responses via Derepressing RELA Expression. mBio 10, e00937–19 (2019).

15. Bravo, J. I. et al. An eQTL-based approach reveals candidate regulators of LINE-1 RNA levels in lymphoblastoid cells. PLoS Genet 20, e1011311 (2024).

16. The GTEx Consortium atlas of genetic regulatory effects across human tissues.

17. Fang, L. et al. The Farm Animal Genotype–Tissue Expression (FarmGTEx) Project. Nat Genet 57, 786–796 (2025).

18. Teng, J. et al. A compendium of genetic regulatory effects across pig tissues. Nat Genet 56, 112–123 (2024).

19. Liu, S. et al. A multi-tissue atlas of regulatory variants in cattle. Nat Genet 54, 1438–1447 (2022).

20. Guan, D. et al. Genetic regulation of gene expression across multiple tissues in chickens. Nat Genet 57, 1298–1308 (2025).

21. Deschamps-Francoeur, G., Simoneau, J. & Scott, M. S. Handling multi-mapped reads in RNA-seq. Computational and Structural Biotechnology Journal 18, 1569–1576 (2020).

22. Arribas, Y. A. et al. Transposable element exonization generates a reservoir of evolving and functional protein isoforms. Cell 187, 7603–7620.e22 (2024).

23. Li, H., Ruan, J. & Durbin, R. Mapping short DNA sequencing reads and calling variants using mapping quality scores. Genome Res. 18, 1851–1858 (2008).

24. Jeong, H.-H., Yalamanchili, H. K., Guo, C., Shulman, J. M. & Liu, Z. An ultra-fast and scalable quantification pipeline for transposable elements from next generation sequencing data. in Biocomputing 2018 168–179 (WORLD SCIENTIFIC, Kohala Coast, Hawaii, USA, 2018). doi:10.1142/9789813235533_0016.

25. Yang, W. R., Ardeljan, D., Pacyna, C. N., Payer, L. M. & Burns, K. H. SQuIRE reveals locus-specific regulation of interspersed repeat expression. Nucleic Acids Research 47, e27–e27 (2019).

26. Jin, Y., Tam, O. H., Paniagua, E. & Hammell, M. TEtranscripts: a package for including transposable elements in differential expression analysis of RNA-seq datasets. Bioinformatics 31, 3593–3599 (2015).

27. Bendall, M. L. et al. Telescope: Characterization of the retrotranscriptome by accurate estimation of transposable element expression. PLoS Comput Biol 15, e1006453 (2019).

28. Ansaloni, F., Gualandi, N., Esposito, M., Gustincich, S. & Sanges, R. TEspeX: consensus-specific quantification of transposable element expression preventing biases from exonized fragments. Bioinformatics 38, 4430–4433 (2022).

29. Li, H. et al. The Sequence Alignment/Map format and SAMtools. Bioinformatics 25, 2078–2079 (2009).

30. Quinlan, A. R. & Hall, I. M. BEDTools: a flexible suite of utilities for comparing genomic features. Bioinformatics 26, 841–842 (2010).

31. Harris, C. R. et al. Array programming with NumPy. Nature 585, 357–362 (2020).

32. Pedregosa, F. et al. Scikit-learn: Machine Learning in Python. MACHINE LEARNING IN PYTHON.

33. Do, C. B. & Batzoglou, S. What is the expectation maximization algorithm? Nat Biotechnol 26, 897–899 (2008).

34. Zhao, M., Liu, D. & Qu, H. Systematic review of next-generation sequencing simulators: computational tools, features and perspectives. Briefings in Functional Genomics elw012 (2016) doi:10.1093/bfgp/elw012.

35. Fairley, S., Lowy-Gallego, E., Perry, E. & Flicek, P. The International Genome Sample Resource (IGSR) collection of open human genomic variation resources. Nucleic Acids Research 48, D941–D947 (2020).

36. Xu, J. et al. A comprehensive benchmarking with interpretation and operational guidance for the hierarchy of topologically associating domains. Nat Commun 15, 4376 (2024).

37. Hermant, C. & Torres-Padilla, M.-E. TFs for TEs: the transcription factor repertoire of mammalian transposable elements. Genes Dev. 35, 22–39 (2021).

38. Burbage, M. et al. Epigenetically controlled tumor antigens derived from splice junctions between exons and transposable elements. Sci. Immunol. 8, eabm6360 (2023).

39. Smit, A. F. A. & Green, P. Using RepeatMasker to Identify Repetitive UNIT 4.10 Elements in Genomic Sequences. Current Protocols in Bioinformatics.

40. Schubert, E., et al. DBSCAN Revisited, Revisited. ACM Transactions on Database Systems 42.

41. Deng, D. DBSCAN Clustering Algorithm Based on Density. in 2020 7th International Forum on Electrical Engineering and Automation (IFEEA) 949–953 (IEEE, Hefei, China, 2020). doi:10.1109/IFEEA51475.2020.00199.

42. Kovaka, S. et al. Transcriptome assembly from long-read RNA-seq alignments with StringTie2. Genome Biol 20, 278 (2019).

43. Tian, X. et al. Widespread impact of transposable elements on the evolution of post-transcriptional regulation in the cotton genus Gossypium. Genome Biol 26, 60 (2025).

44. Teng, J., et al. OmiGA: A Toolkit for Ultra-efficient Molecular Trait Analysis in Complex Populations.

45. Storey, J. D. A Direct Approach to False Discovery Rates. Journal of the Royal Statistical Society Series B: Statistical Methodology 64, 479–498 (2002).

46. Cingolani, P. et al. A program for annotating and predicting the effects of single nucleotide polymorphisms, SnpEff: SNPs in the genome of Drosophila melanogaster strain w^1118^LJ; iso-2; iso-3. Fly 6, 80–92 (2012).

47. Mountjoy, E. et al. An open approach to systematically prioritize causal variants and genes at all published human GWAS trait-associated loci. Nat Genet 53, 1527–1533 (2021).

48. Da, Y. A Large-Scale Genome-Wide Association Study in U.S. Holstein Cattle.

49. Guan, D. et al. Genetic regulation of gene expression across multiple tissues in chickens. Nat Genet 57, 1298–1308 (2025).

50. Gusev, A. et al. Integrative approaches for large-scale transcriptome-wide association studies. Nat Genet 48, 245–252 (2016).

51. Tabrizi, S. J., Flower, M. D., Ross, C. A. & Wild, E. J. Huntington disease: new insights into molecular pathogenesis and therapeutic opportunities. Nat Rev Neurol 16, 529–546 (2020).

52. Ortíz-Fernández, L., Martín, J. & Alarcón-Riquelme, M. E. A Summary on the Genetics of Systemic Lupus Erythematosus, Rheumatoid Arthritis, Systemic Sclerosis, and Sjögren’s Syndrome. Clinic Rev Allerg Immunol 64, 392–411 (2022).

53. Beydon, M. et al. Epidemiology of Sjögren syndrome. Nat Rev Rheumatol 20, 158–169 (2024).

54. Niewold, T. B. et al. Association of the IRF5 risk haplotype with high serum interferon-α activity in systemic lupus erythematosus patients. Arthritis & Rheumatism 58, 2481–2487 (2008).

55. Ban, T., Sato, G. R. & Tamura, T. Regulation and role of the transcription factor IRF5 in innate immune responses and systemic lupus erythematosus. International Immunology 30, 529–536 (2018).

56. Li, D., et al. IRF5 genetic risk variants drive myeloid-specific IRF5 hyperactivation and presymptomatic SLE. JCI Insight 5, e124020 (2020).

57. Lu, M. et al. TWAS Atlas: a curated knowledgebase of transcriptome-wide association studies. Nucleic Acids Research 51, D1179–D1187 (2023).

58. Liu, M., Wang, S., Liang, Y., Fan, Y. & Wang, W. Genetic polymorphisms in genes involved in the type I interferon system (STAT4 and IRF5): association with Asian SLE patients. Clin Rheumatol 43, 2403–2416 (2024).

59. Hubley, R., Wheeler, T. J. & Smit, A. F. A. Accuracy of multiple sequence alignment methods in the reconstruction of transposable element families. NAR Genomics and Bioinformatics 4, lqac040 (2022).

60. Brown, A. A. et al. Genetic analysis of blood molecular phenotypes reveals common properties in the regulatory networks affecting complex traits. Nat Commun 14, 5062 (2023).

61. Jurka, J. et al. Repbase Update, a database of eukaryotic repetitive elements. Cytogenet Genome Res 110, 462–467 (2005).

62. Altschul, S. Gapped BLAST and PSI-BLAST: a new generation of protein database search programs. Nucleic Acids Research 25, 3389–3402 (1997).

63. Zdobnov, E. M., Lopez, R., Apweiler, R. & Etzold, T. The EBI SRS server—recent developments. Bioinformatics 18, 368–373 (2002).

64. Barrett, T. et al. NCBI GEO: mining tens of millions of expression profiles--database and tools update. Nucleic Acids Research 35, D760–D765 (2007).

65. Katz, K. et al. The Sequence Read Archive: a decade more of explosive growth. Nucleic Acids Research 50, D387–D390 (2022).

66. Chen, S. Ultrafast one-pass FASTQ data preprocessing, quality control, and deduplication using fastp. iMeta 2, e107 (2023).

67. Dobin, A. et al. STAR: ultrafast universal RNA-seq aligner. Bioinformatics 29, 15–21 (2013).

68. Liao, Y., Smyth, G. K. & Shi, W. featureCounts: an efficient general purpose program for assigning sequence reads to genomic features. Bioinformatics 30, 923–930 (2014).

69. Ihaka, R. & Gentleman, R. R: A Language for Data Analysis and Graphics. Journal of Computational and Graphical Statistics 5, 299–314 (1996).

70. Gay, N. R. et al. Impact of admixture and ancestry on eQTL analysis and GWAS colocalization in GTEx. Genome Biol 21, 233 (2020).

71. Robinson, M. D., McCarthy, D. J. & Smyth, G. K. edgeRLJ: a Bioconductor package for differential expression analysis of digital gene expression data. Bioinformatics 26, 139–140 (2010).

72. Raivo Kolde. pheatmap: Pretty Heatmaps. 1.0.12 10.32614/CRAN.package.pheatmap (2010).

73. Yin, L., et al. rMVP: A Memory-Efficient, Visualization-Enhanced, and Parallel-Accelerated Tool for Genome-Wide Association Study. Genomics, Proteomics & Bioinformatics 19, 619–628 (2021).

74. Wilkinson, L. ggplot2: Elegant Graphics for Data Analysis by WICKHAM, H. Biometrics 67, 678–679 (2011).

